# A portable library to support the SBML Layout Extension

**DOI:** 10.1101/035725

**Authors:** J. Kyle Medley, Kiri Choi, Herbert M. Sauro

## Abstract

The SBML layout extension enables SBML models to encode layout information which describes the graphical depiction of model elements. In this application note, we describe libSBNW, a portable library that supports the SBML layout extension and can automatically generate layout for SBML models. The library can be used to automatically generate layout information for SBML models lacking it, or to edit coordinate information already encoded in a model. We provide C and Python APIs to allow other applications to host the library or to use it directly from the Python console. We show that the library is sufficient for creating a graphical application for displaying and editing layout information. The library is open-source and licensed under the BSD 3-clause license. Project source code, downloads, documentation and binaries for Windows and Mac OS X are available at https://github.com/sys-bio/sbnw. The library is also included in Tellurium, available at http://tellurium.analogmachine.org/. Video tutorials are available at http://0u812.github.io/sbnw/tutorials/.

## 2 Introduction

SBML [Hucka et al., 2003] is the de facto standard for exchanging biochemical network models (Sauro, 2014). The SBML effort has spawned a great variety of extensions and other formats [Dräger et al., 2014]. One such extension is the SBML layout extension [Gauges et al., 2006], which is embedded in an SBML document. The extension allows software to describe the graphical layout of a biochemical network in terms of species, reactions, compartments and modifiers. On the other hand, the layout extension does not provide information on the detailed rendering of the network, i.e. colors or shapes of symbols. Such details are reserved for the render extension [Gauges, 2009].

However, many SBML models contain no layout information [Bergman and Sauro, 2006], and this is a barrier to developing graphical software tools for interacting with SBML. Therefore, it is desirable to have a library for assigning layout information to SBML models created before the introduction of the layout extension, and to provide an easy way to encode layout information in future SBML models.

In this application note we present an updated library based on the original SBW layout engine [Deckard et al., 2006]. The original library was written in C# and was tailored specifically for use by SBW. Here, we describe a new version, written in C/C+++, with fewer dependencies, Python bindings, sample applications and additional enhancements.

## 3 Methods

**SBML Layout Extension Support** The library is designed for compatibility across SBML level 2/3, and uses libSBML for reading and writing content [Bornstein et al., 2008]. The SBML layout extension is used to store visual layout information. If the input model does not have a layout, one can be automatically generated by the library.

**Autolayout Algorithm** The library automatically generates layout information encoding node and reaction centroid coordinates using the Fruchterman-Reingold (FR) algorithm, which has been shown to be robust in the face of variegated graph topologies and faithfully reproduces the underlying symmetry [Fruchterman and Reingold, 1991].

**Selectively lock Nodes** Users can specify one or more nodes to be locked when the layout algorithm is executed. This ensures that the positions of these nodes do not change when the layout algorithm is applied. Node locking is particularly useful for users who want to fine-tune the automatic layout process.

**Support for Alias Nodes** Models with a high degree of connectivity can be problematic for visualization methods due to the overlap between edges that occurs when the underlying graph is embedded in 2D space. We solve this problem by providing the user with the ability to create *alias nodes*. In general, any node of degree n can be decomposed into n alias nodes, each of which is rendered separately. Using this method, we can reduce the connectivity of any network graph. The FR-algorithm is particularly adept at laying out such reduced graphs without overlapping edges [Fruchterman and Reingold, 1991], enabling lucid visualization of complex networks.

**Export of Rendered Model** A rendered model may be exported to a raster format (Portable Network Graphics) or one of several vector formats (Scalable. Vector Graphics, TikZ).

**Supply Ancillary graphical Information** Visual features such as arrowheads/endcaps are supplied by the library, and may be used to render network diagrams as shown in Figure 1. Five endcap styles are supported and are tied to species roles in SBML reactions: Substrate, product, modifier, activator and inhibitor.

**Figure 1:**
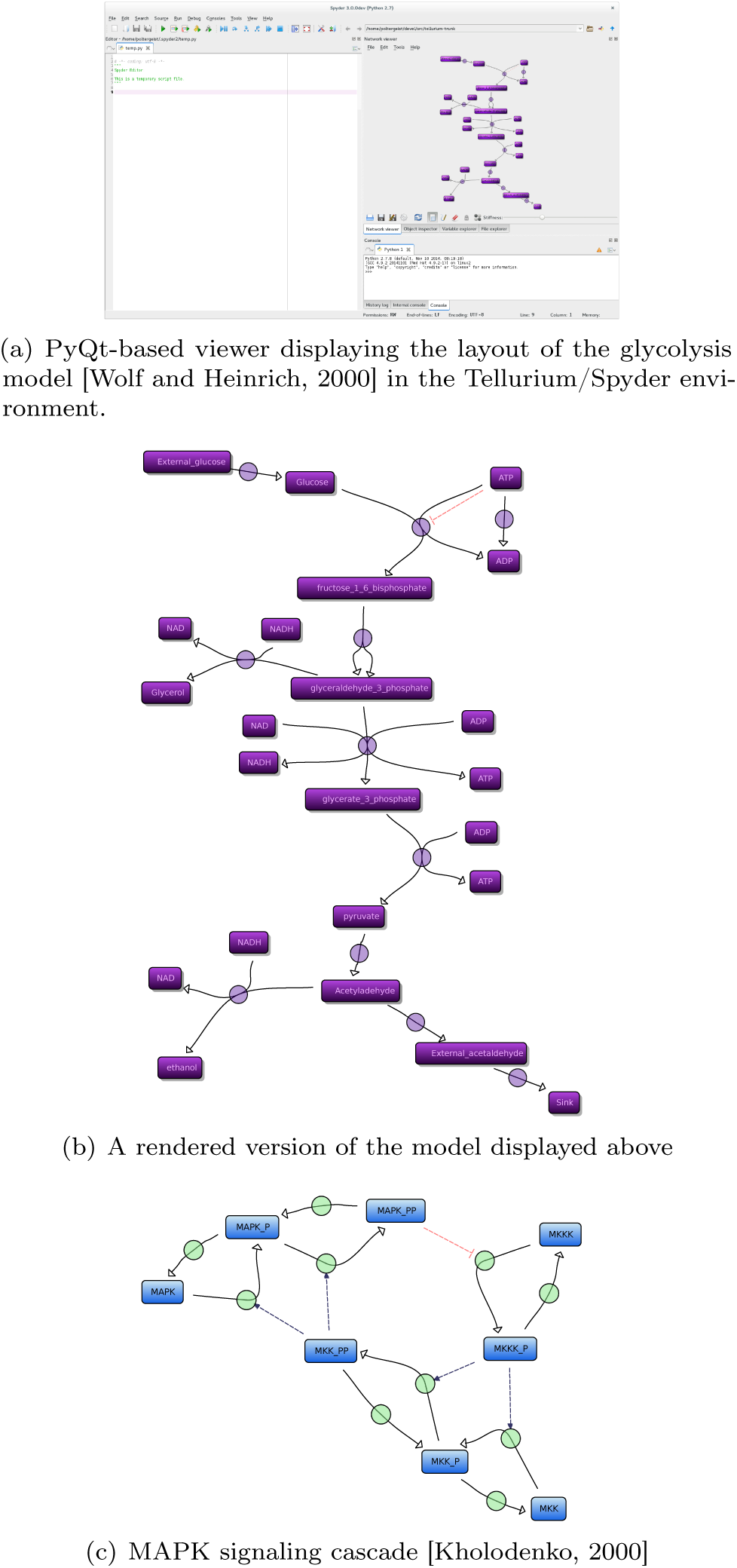
Demonstration of the Python-based network viewer

**Language Bindings** Public APIs for C and Python are provided. For example, the following Python code loads an SBML model, applies the FR autolayout algorithm to the network, and saves the result as output.xml in the current directory:

~~~
# import the sbnw Python module
import sbnw
# load the model
model = sbnw. loadsbml (’ model. xml ’)
# seed node coordinates randomly
model.network.randomize()
# apply the FR-algorithm
model.network.autolayout ()
# Save new SBML to file
model.save( ’output.xml ’)
~~~

## 4 Results

We have deployed libSBNW in the context of the Tellurium biological modeling environment (http://tellurium.analog-machine.org/) using the Tellurium’s Spyder-based plugin architecture. Tellurium includes the nwed (network editor) Python module, which may be used to communicate with the layout plugin, and the SBML model may be set or retrieved via nwed.setsbml and nwed.getsbml respectively.

~~~
sbmlStr = ’’’ … ’’’
import nwed
nwed.setsbml(sbmlSstr)
~~~

## Acknowledgments

We acknowledge Stanley Gu for his shared web development expertise, Lucian Smith for his work on the Antimony language, Jonathan Gallimore for his work on the C API, Andy Somogyi for his shared expertise in Python extension modules, and Kaylene Stocking for testing the network viewer.

## Funding

The authors are most grateful to generous funding from the National Institute of General Medical Sciences of the National Institutes of Health under award R01-GM081070. The content is solely the responsibility of the authors and does not necessarily represent the official views of the National Institutes of Health.

